# Increased activity of PRMT5-MEP50 complex improves survival of chromosomally unstable cancer cells by increasing tolerance to protein aggregation and proteotoxicity

**DOI:** 10.1101/2025.09.12.675799

**Authors:** Maria F. Suarez Peredo Rodriguez, Laura J. Jilderda, Catalina Gaviria Agudelo, Judith E. Simon, Alexander M. Heberle, Alex van Kaam, Petra L. Bakker, Christy Hong, Maurits Roorda, Genevieve Lavoie, Philippe P. Roux, Andrea Oeckinghaus, Marcel A.T.M. van Vugt, Kathrin Thedieck, Floris Foijer

**Affiliations:** European Research Institute for the Biology of Ageing (ERIBA), University of Groningen, University Medical Centre Groningen, 9713 AV Groningen, The Netherlands; Institute of Biochemistry and Center for Molecular Biosciences Innsbruck, University of Innsbruck, Innsbruck 6020, Austria; Department of Medical Oncology, University of Groningen, University Medical Centre Groningen, 9713 AV Groningen, The Netherlands; Institute for Research in Immunology and Cancer (IRIC), Université de Montréal, Montreal, Canada; Department of Pathology and Cell Biology, Faculty of Medicine, Université de Montréal, Montreal, Quebec, Canada; Department Metabolism, Senescence and Autophagy, Research Center One Health Ruhr, University Alliance Ruhr & University Hospital Essen, University Duisburg-Essen, Essen, Germany; Center of Medical Biotechnology, Faculty of Biology, University of Duisburg-Essen, 45141 Essen, Germany; German Cancer Consortium (DKTK), partner site Essen/Duesseldorf, a partnership between German Cancer Research Center (DKFZ) and University Hospital Essen, 45147 Essen, Germany; Laboratory of Pediatrics, Section Systems Medicine of Metabolism and Signaling, University of Groningen, University Medical Center Groningen, 9713 GZ Groningen, The Netherlands; Freiburg Materials Research Center FMF, Albert-Ludwigs-University of Freiburg,79104 Freiburg, Germany

**Keywords:** cancer, aneuploidy, chromosomal instability, protein aggregation, proteotoxicity, proliferation, PRMT5, MEP50

## Abstract

Most cancers display chromosomal instability (CIN), a condition that leads to increased rates of chromosome missegregation and thus yields aneuploidy. CIN and aneuploidy are detrimental to healthy cells and therefore, aneuploid cells rely on aneuploidy-tolerating mechanisms to adopt a malignant fate. We previously found PRMT5 to be frequently amplified in a mouse model for aneuploid T cell lymphoblastic lymphoma. In this study, we investigated a possible role of PRMT5 as an aneuploidy tolerating gene. We report that PRMT5 is prone to aggregation when expressed at supra-stoichiometric levels compared to its obligate partner protein MEP50 (methylosome protein 50, WDR77). Intriguingly, we also find that protein aggregation, induced by CIN, is mitigated by jointly increased expression of PRMT5 and MEP50. Accordingly, concomitant PRMT5:MEP50 expression renders cancer cells less sensitive to proteasome inhibitors and CIN while inhibition sensitizes cells to CIN. Our findings provide a possible explanation for why PRMT5 and MEP50 display increased expression, particularly in aneuploid cancers and might reveal a targetable vulnerability of aneuploid cancer.

## Introduction

All somatic cells in the human body contain two sets of chromosomes (chr.), with each set of 23 chromosomes inherited from one parent. Upon cell division, chromosomes are replicated and need to be evenly distributed over the two emerging daughter cells to maintain correct chromosome numbers. Errors during chromosome segregation, a condition known as chromosomal instability (CIN), lead to cells with abnormal chromosome numbers, a state defined as aneuploidy^1^.

CIN and aneuploidy impair proliferation across cell types and organisms from yeast^2^ to mammalian cells^3–5^ Aneuploidy compromises cellular fitness as it results in an altered cellular metabolism, leads to an increased production of reactive oxygen species (ROS)^6^ and yields unbalanced protein expression from the aneuploid chromosomes^7^. Imbalanced protein complex stoichiometry leads to protein misfolding and aggregation ^2,8–12^, which results in an increased sensitivity to compounds that interfere with proteostasis^8,13^.

Despite their detrimental effects on untransformed cells, aneuploidy and CIN are common to cancer cells, which are characterized by uncontrolled cell proliferation^14,15^. More than 80% of all solid tumors are aneuploid, and CIN has been associated with increased metastatic potential, increased rates of tumor recurrence, an increased frequency of drug resistance, and thus, overall, a poor prognosis^16–19^. At the cellular level, CIN promotes karyotype evolution, reshuffling chromosomes carrying oncogenes or tumor suppressor genes, to yield a new optimized balance between these two types of genes, thus driving new cancer prone karyotypes and cancer cell selection^20,21^. Conversely, chromosome copy number changes also alter expression of many non-cancer-related (passenger) genes, some of which decrease cellular fitness and trigger a cellular stress response^20^. This implies that premalignant cells might need to overcome CIN-imposed stresses to become tumorigenic and furthermore that such coping mechanisms might provide targetable vulnerabilities of CIN^HIGH^ cancers.

We previously identified PRMT5 as a potential driver of CIN^HIGH^ murine T-cell acute lymphoblastic lymphoma (T-ALL). PRMT5 is located on chromosome 14, which we found commonly amplified in these T-ALLs. In line, PRMT5 expression was consistently upregulated in these T-ALLs^22,23^. Indeed, PRMT5 expression is frequently increased in many cancer types, including lymphoma^24^ and is therefore considered to be a potential clinical target to treat cancer, evidenced by several clinical trials of PRMT5 inhibitors in hematologic and solid malignancies^25^. PRMT5 is a class II protein arginine methyltransferase that, together with its obligate binding partner and adaptor protein MEP50^26^, symmetrically di-methylates arginine residues^27–30^. The PRMT5:MEP50 complex methylates histone tails as well as a plethora of other substrates that control transcription, mRNA processing and translation, proteostasis, DNA repair, cell cycle progression, and oncogenic signaling^24^. Depletion of PRMT5 is embryonic lethal and leads to death of most cell types, attesting to its essential function^28^.

As PRMT5 is frequently overexpressed in various types of cancer and recurrently gained in mouse models for CIN^HIGH^ T-ALL, we hypothesized that PRMT5 contributes to aneuploidy tolerization. We therefore studied the effect of controlled PRMT5 overexpression in isogenic CIN^LOW^ and CIN^HIGH^ cell models. We report that PRMT5 is prone to aggregation when expressed without its obligate adaptor MEP50, particularly in CIN^HIGH^ cells. However, when PRMT5 and MEP50 are co-induced to similar levels, protein aggregation is suppressed, both in CIN^LOW^ and CIN^HIGH^ cells. In line with this, PRMT5:MEP50 co-overexpressing cells are desensitized to the CIN-inducing compounds reversine^31^ and cpd-5^32^, as well as the proteasome inhibitor MG132, whereas PRMT5 expression alone renders cells sensitive to these compounds. Furthermore, concerted PRMT5:MEP50 induction, but not PRMT5 induction alone, prevent aggregation of a huntingtin-based probe. Our observations are conserved across cancer types as high PRMT5 expression positively correlates with MEP50 levels, aneuploidy scores and resistance to proteasome inhibitors in DepMap data. Altogether, our observations suggest that a stoichiometrically increased PRMT5-MEP50 complex improves fitness of CIN^HIGH^ cells by buffering the proteotoxic effects of CIN-induced protein aggregation. Conversely, induced expression of PRMT5 alone enhances proteoxicity and is selected against. Therefore, disturbance of the MEP50:PRMT5 equilibrium may offer a new targetable vulnerability of cancer cells that exhibit CIN.

## Results

### Increased expression of PRMT5 leads to its accumulation in cytoplasmic foci that increase in number following the induction of CIN

To investigate the function of PRMT5 in cells with CIN, we first increased PRMT5 expression in otherwise unperturbed RPE1 and MCF10A cells, both near-diploid, non-transformed human cell lines^33,34^. We engineered doxycycline-inducible^35^ GFP-PRMT5 and PRMT5-GFP retroviral vectors to control PRMT5 expression levels and monitored its localization. We titrated expression to match a cancer-physiologically relevant range, ∼2-6-fold higher than in control cells^22^ (**Fig. 1A**, **B** and **Supp.** Fig. 1A). Both GFP-PRMT5 and PRMT5-GFP migrated at the expected height (∼100 kDa) with a minor fraction of the fusion proteins being degraded. Live cell imaging revealed that in both cell types the two GFP fusion proteins accumulated in cytoplasmic foci (**Fig. 1C, D**, **Supp.** Fig. 1B). This was confirmed by immunofluorescence (IF) using two different PRMT5 antibodies (**Fig. 1E**). To rule out that foci formation was caused by GFP-fusion, we engineered HA-tagged PRMT5, which also localized to cytoplasmic foci (**Supp.** Fig. 1C).

**Figure 1.**
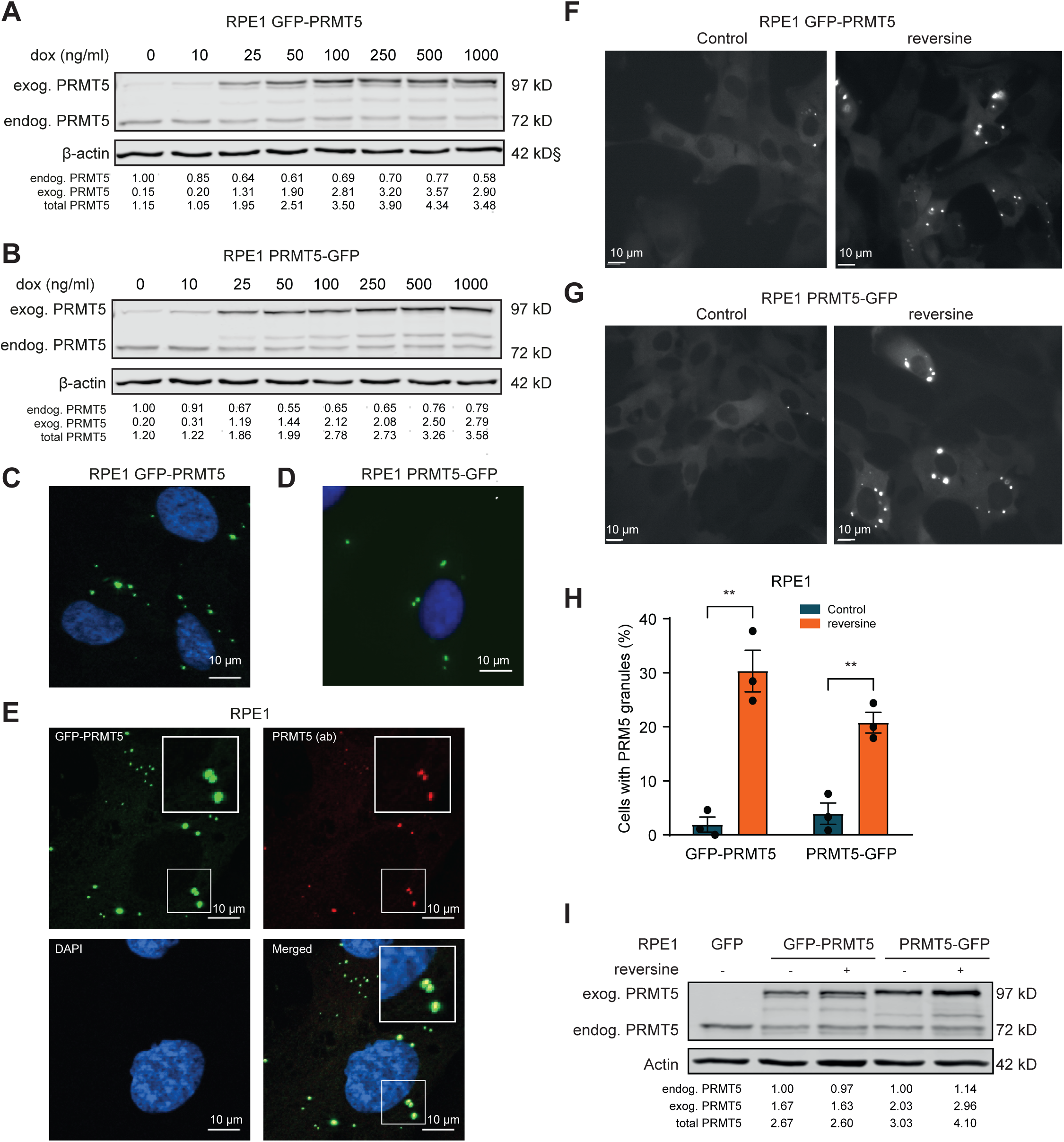
Overexpressed PRMT5 accumulates in cytoplasmic foci and aneuploidization increases the number of PRMT5 foci. (A-B) Immunoblots showing PRMT5 and β-actin expression in **(A)** RPE1 rtTA GFP-PRMT5 and **(B)** RPE1 rtTA PRMT5-GFP cells treated for 48 h with 0-1000 ng/ml dox. Quantification of PRMT5 bands is shown under each blot. **(C-D)** Representative stills of live cell immunofluorescence microscopy detection of GFP-tagged PRMT5 and nuclei/DAPI in RPE1 rtTA GFP-PRMT5 and RPE1 rtTA PRMT5-GFP cells treated for 24 h with 100 ng/ml dox. **(E)** Detection of PRMT5 by immunofluorescence microscopy in RPE1 rtTA GFP-PRMT5 RPE1 cells treated for 24 h with 500 ng/ml dox and stained with a PRMT5 antibody (Bethyl, A300-850A). Scale bar = 10 μm. Figure represents one experiment. Representative stills of live-cell imaging of (**F**) RPE1 rtTA GFP-PRMT5 cells and **(G)** RPE1 rtTA PRMT5-GFP cells treated for 48 h with 500 nM rev and 24 h with 25 ng/ml dox prior to imaging. Scale bar = 10 μm. **(H)** Quantification of live cell imaging shown in **(F,G)** For statistical analysis, a two-tailed student’s t-test was applied: **, p ≤ 0.01. Data represent 3 biological replicates. Data are presented as mean ± SEM. **(I)** Western blot showing PRMT5 and β-actin expression in RPE1 rtTA GFP-PRMT5 and PRMT5-GFP cells treated for 72 h with 500 nM rev and for 48 h with 25 ng/mL dox. Quantification relative to endogenous PRMT5 in the control condition is given below the blot panels.

Next, we induced CIN in these cell lines using the serine/threonine-protein kinase Mps1 inhibitor reversine (rev)^31^. To determine the effect of CIN on PRMT5 foci formation, we titrated PRMT5 expression levels so that only 2-5% cells displayed PRMT5 foci in the control condition (25 ng/ml) (**Fig. 1F, G**; left panels). Rev increased the fraction of cells with PRMT5 granules by ∼4-10 fold (**Fig. 1F-H**), despite comparable PRMT5 levels in control and rev-treated cells (**Fig. 1I**). Similar results were obtained for MCF10A cells (**Sup.** Fig. 1 **D, E**; ∼two-fold increase in foci). This suggests that CIN promotes the formation of PRMT5 granules when its expression is increased.

PRMT5 methylates the core stress granule (SG) component G3BP1^36–39^ and PRMTs have been shown to control SG assembly^40–42^. We therefore investigated whether the PRMT5 foci were SGs. GFP-PRMT5 foci did not co-localize with G3BP1. In fact, no G3BP1 granules were observable in GFP-PRMT5 expressing RPE1 cells (**Fig. 2A**, upper panel). We treated cells with arsinite to induce SGs^42^ but also in this setting we failed to detect colocalization between mCherry-tagged G3BP1^21^ and GFP-PRMT5 (**Fig. 2A**, middle and lower panels). This demonstrates that GFP-PRMT5 foci are not SGs.

**Figure 2.**
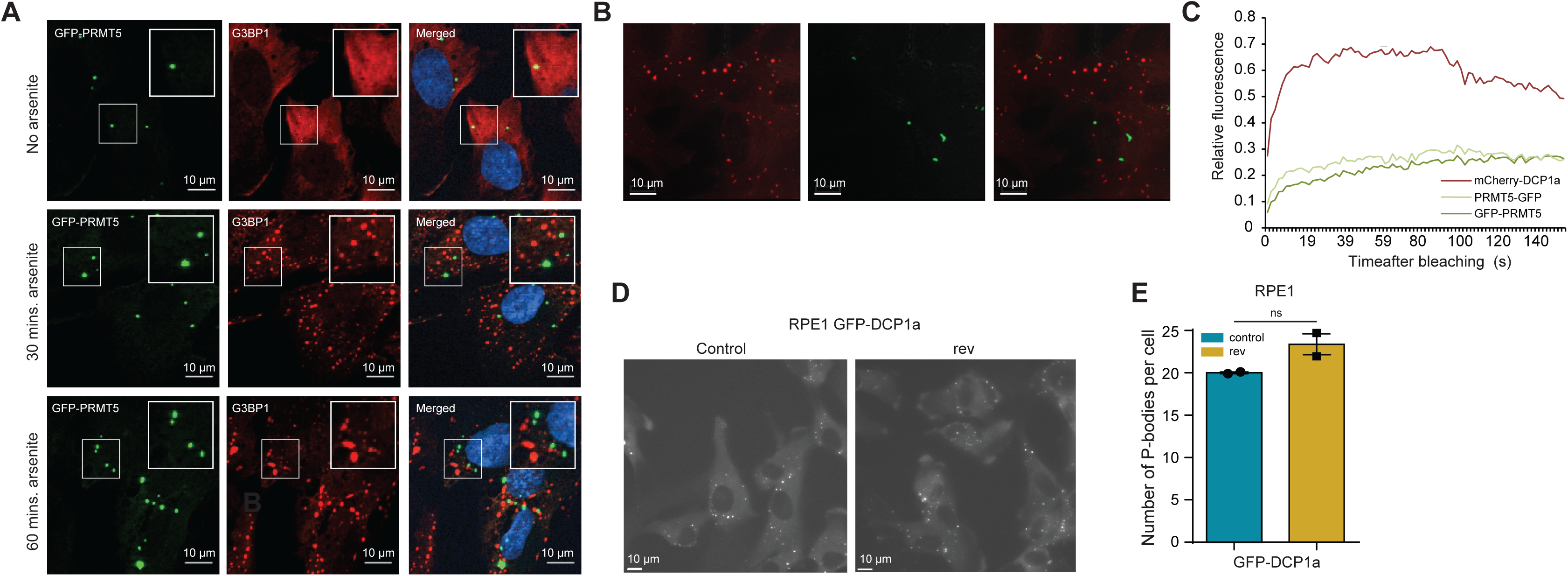
Overexpressed PRMT5 does not co-localize with p-bodies or stress granules. **(A)** Immunofluorescence detection of GFP-PRMT5, G3BP1, and nuclei/DAPI in RPE1 rtTA GFP-PRMT5 cells treated with 500ng/ml dox for 24 h in and then stimulated with 500 µM arsinite for 30 or 60 min. Scale bar = 10 µm. **(B)** Representative stills of live-cell imaging of RPE1 rtTA PRMT5 GFP + mCherry-DCP1A cells treated for 48 h with 500 ng/mL dox prior to imaging. Scale bar = 10 µm. **(C)** FRAP analysis of RPE1 rtTA GFP-PRMT5, PRMT5-GFP, or mCherry-DCP1A cells. Prior to imaging, cells were treated for 48 h with 500 ng/ml dox. y-axis represents recovery and x-axis time (s) after bleaching. Data represent one experiment. **(D)** Representative stills of live-cell imaging of RPE1 rtTA GFP-DCP1A prior to imaging treated for 24h with 500 ng/ml dox (left) and 500 ng/ml plus 500 nM rev (right). Scale bar is 10 μm. **(E)** Quantification of p-bodies in live-cell imaging shown in **(D)**. For statistical analysis, a two-tailed student’s t-test was applied: ns, nonsignificant. Data represent 3 biological replicates and indicate the mean ± SEM.

PRMT5 has been linked with p-bodies (PB), the centers of mRNA turnover^43^. To test if the PRMT5 foci we observed were PBs, we expressed an PB reporter, mCherry-tagged-DCP1A^44^, together with GFP-PRMT5, but also here we failed to detect co-localization (**Fig. 2B**). Fluorescence recovery after photobleaching (FRAP) microscopy revealed that mCherry-DCP1A fluorescence recovered rapidly, confirming a high turnover rate in PBs^45^, whereas GFP-PRMT5 fluorescence did not recover within the timeframe of the experiment (**Fig. 2C**). We also compared PB numbers (mCherry-DCP1a-foci) between cells with and without induced CIN and found no difference (**Fig. 2D, E**), reinforcing that PRMT5 foci are not PBs. We conclude that increased expression of PRMT5 leads to the formation of static cytoplasmic foci, that are increased in number when cells exhibit CIN and that are neither SGs nor PBs.

### PRMT5 foci lack its obligate binding partner MEP50

To gain insight into the composition of the PRMT5 foci, we analyzed the proteins in GFP-PRMT5 foci under rev and control conditions using label-free mass spectrometry-based proteomics (**Fig. 3A**). GFP-PRMT5 foci were isolated from cell lysates by centrifugation, followed by immunoprecipitation (IP) of GFP. GFP-PRMT5 foci showed enrichment for proteins related to RNA processing, without overt differences between mock- and rev-treated cells (**Fig. 3B**). This indicates that the composition of PRMT5 foci is not affected by CIN. We also determined binding partners of GFP-PRMT5 outside of PRMT5 foci (i.e. binding partners in the supernatant fraction of the centrifuged cell lysates) and found many interactions to be overlapping with interactors of GFP-PRMT5 present in PMRT5 foci. However, several proteins known to be required for PRMT5 function were only found to bind to GFP-PRMT5 when not in foci, including the PRMT5 binding partner MEP50^46,47^, and RioK1 (Rio kinase 1), one of the adaptor proteins that determine PRMT5 substrate specificity in the methylosome complex^48^ (**Fig. 3C, D**).

**Fig 3.**
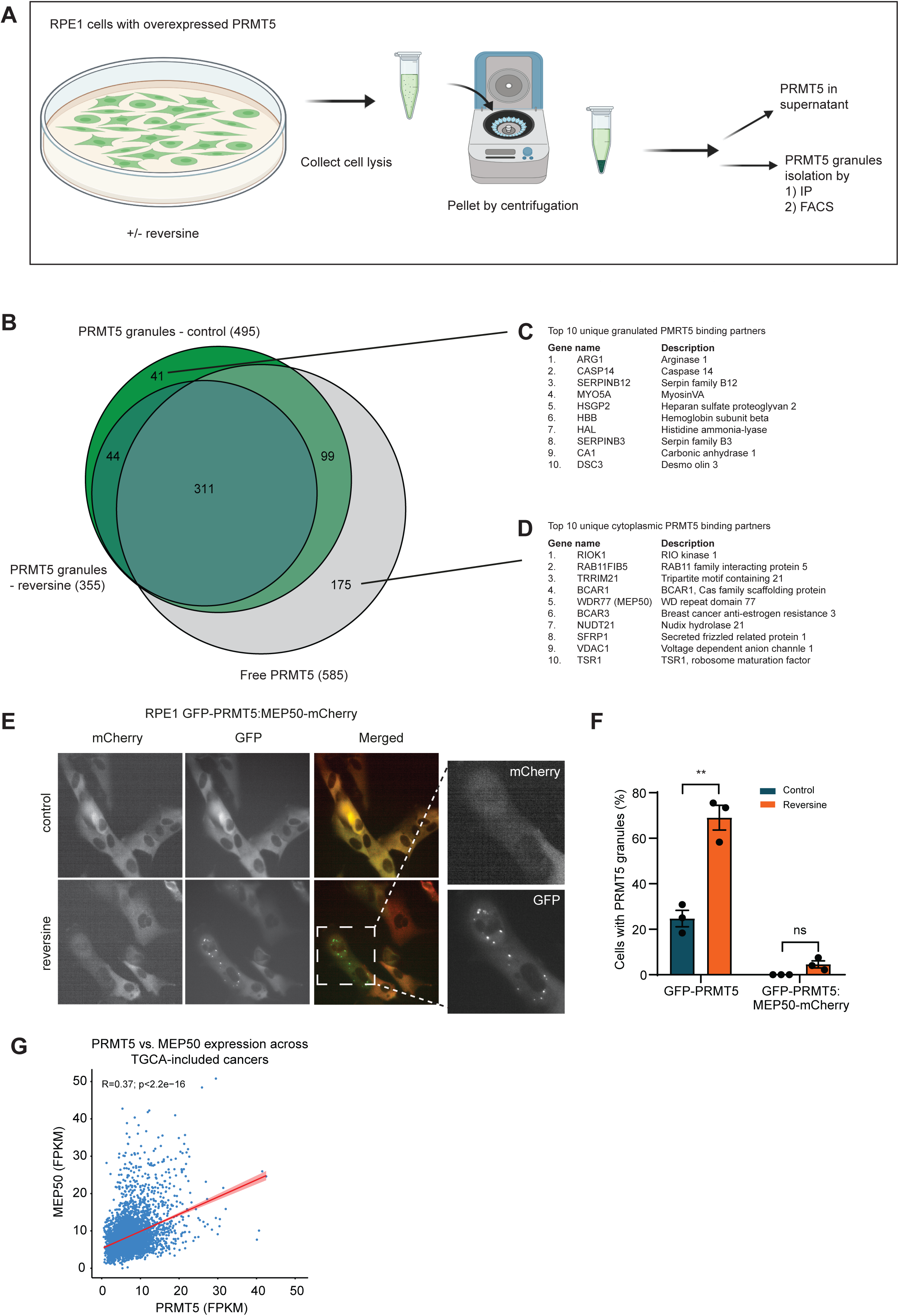
Stoichiometric imbalances of the PRMT5-MEP50 complex lead to PRMT5-foci formation. **(A)** Experimental overview for mass spectrometry experiment. Free PRMT5 and PRMT5 foci were isolated using immunoprecipitation or FACS from RPE1 expressing GFP-PRMT5 or PRMT5-GFP cells treated carrier control or rev. The proteins present in the different fractions were then analyzed using label-free mass spectrometry. **(B)** Venn diagram depicting number of proteins identified in the three fractions: 1) free, cytosolic PRMT5, 2) cytosolic PRMT5 foci induced by 72 h of dox treatment, and 3) cytosolic PRMT5 foci induced by 72 h of dox and 500 nM rev treatment. **(C)** Table showing top 10 unique proteins identified in PRMT5 foci. **(D)** Table showing top 10 unique proteins identified in the free PRMT5 fraction. **(E)** Representative stills of live-cell imaging of RPE1 rtTA GFP-PRMT5-T2A-MEP50-mCherry treated for 48 h with 500 nM rev and for 24 h with 1000 ng/ml dox prior to imaging. **(F)** Quantification of cells with PRMT5 foci of IncuCyte-derived images of RPE1 rtTA GFP-PRMT5 and rtTA GFP-PRMT5-T2A-MEP50-mCherry cells treated with 500 ng/ml dox and 500 nM rev for 7 days. n=3. For statistical analysis, a two-tailed student’s t-test was applied: * p ≤ 0.05, ** p ≤ 0.01, n.s: non-significant. Data are represented as mean ± SEM. **(G)** Correlation plot for PRMT5 and MEP expression across TGCA-included cancer samples.

As MEP50 is a critical cofactor for PRMT5 methyl transferase activity^49^ we tested whether stoichiometrically matched expression of PRMT5 and MEP50 affected PRMT5 foci formation. We engineered RPE1 and MCF10A cells to co-express PRMT5 and MEP50 in a 1:1 stoichiometric ratio using a dox-inducible GFP-PRMT5-T2A-MEP50-mCherry construct. This allowed to induce expression of both proteins by approximately two-fold (**Supp.** Fig. 1F, G). As a control, we examined cells expressing PRMT5 alone. While PRMT5 expression alone yielded PRMT5 foci, their formation was prevented when MEP50 and PRMT5 were expressed concomitantly (**Fig. 3E, F**). Also, upon rev-induced CIN, co-expression of MEP50 largely prevented PRMT5 foci formation (**Fig. 3E, F**). A few cells still showed PRMT5 foci, but these had very low MEP50-mCherry expression (**Fig. 3E**, zoomed), further suggesting that PRMT5 foci formation occurs when PRMT5 is present in supra-stoichiometric levels compared to its binding partner MEP50. If PRMT5 foci compromise cell fitness, one would expect a selective pressure towards coordinated stoichiometric expression of PRMT5 and MEP50. Indeed, PRMT5 and MEP50 expression strongly correlated across expression data from TGCA-included cancers (**3G**). Hence, cancers do not only increase expression of PRMT5 and MEP50 but they also maintain their correct stoichiometry.

### Coordinated MEP50:PRMT5 expression reduces CIN-induced protein aggregation

Aggregation of supra-stoichiometric proteins is common in cells with CIN^50,51^. This aligns with our observation that disturbance of the stoichiometric ratio between PRMT5 and MEP50 leads to the formation of PRMT5 foci. However, this does not explain why rev-induced CIN further promoted PRMT5 foci formation, as the PRMT5 levels were not different to the carrier control (**Fig. 1I**). Also, the near complete suppression of rev-induced PRMT5 foci by MEP50 (**Fig. 3F**) suggested that a reason beyond a disturbed PRMT5:MEP50 stoichiometry accounted for the increased PRMT5 foci numbers in CIN cells. This could imply that MEP50:PRMT5 activity protects cells from CIN-induced protein aggregation. We tested whether expression of PRMT5:MEP50 versus PRMT5 alone affected protein aggregation in a CIN background. Towards this aim, we engineered inducible HA-PRMT5 and HA-PRMT5:MEP50 MCF7 breast cancer cells (**Supp.** Fig. 1H, I) that also expressed a GFP-tagged aggregation-prone protein fragment based on exon 1 of the Huntingtin gene, which contains a stretch of 71 glutamines (HTTEX1-Q71; PolyQ stretch^52^. We compared the number of GFP-PolyQ aggregates between control-and rev-treated MCF7 cells with PRMT5 alone versus PRMT5:MEP50 by time-lapse imaging for up to 20 hours (**Fig. 4**). Control cells without induced PRMT5 or PRMT5:MEP50 expression showed very few PolyQ aggregates (**Fig. 4A, B**, blue and green lines in B, respectively). Rev increased accumulation of PolyQ aggregates (**Fig. 4C, D** blue and green lines in C, respectively), indicating that CIN exacerbated protein aggregation. Induction of PRMT5 alone did not prevent PolyQ aggregation, instead the number of PolyQ aggregates further increased (**Fig. 4C, E**, red line in C). This is in line with our finding that CIN exacerbates PRMT5 foci (**Fig. 1 F-H**) and supports the idea that CIN induces the number of PRMT5 foci because of overall enhanced protein aggregation. PRMT5:MEP50 co-expression had the opposite effect and suppressed CIN-induced PolyQ aggregation (**Fig. 4C, F**, purple line in C). Together, the results suggest that PRMT5 expression alone exacerbates CIN-induced aggregation, whereas stoichiometric PRMT5:MEP50 expression prevents protein aggregation, thus reducing aneuploidy-induced proteotoxic stress.

**Figure 4.**
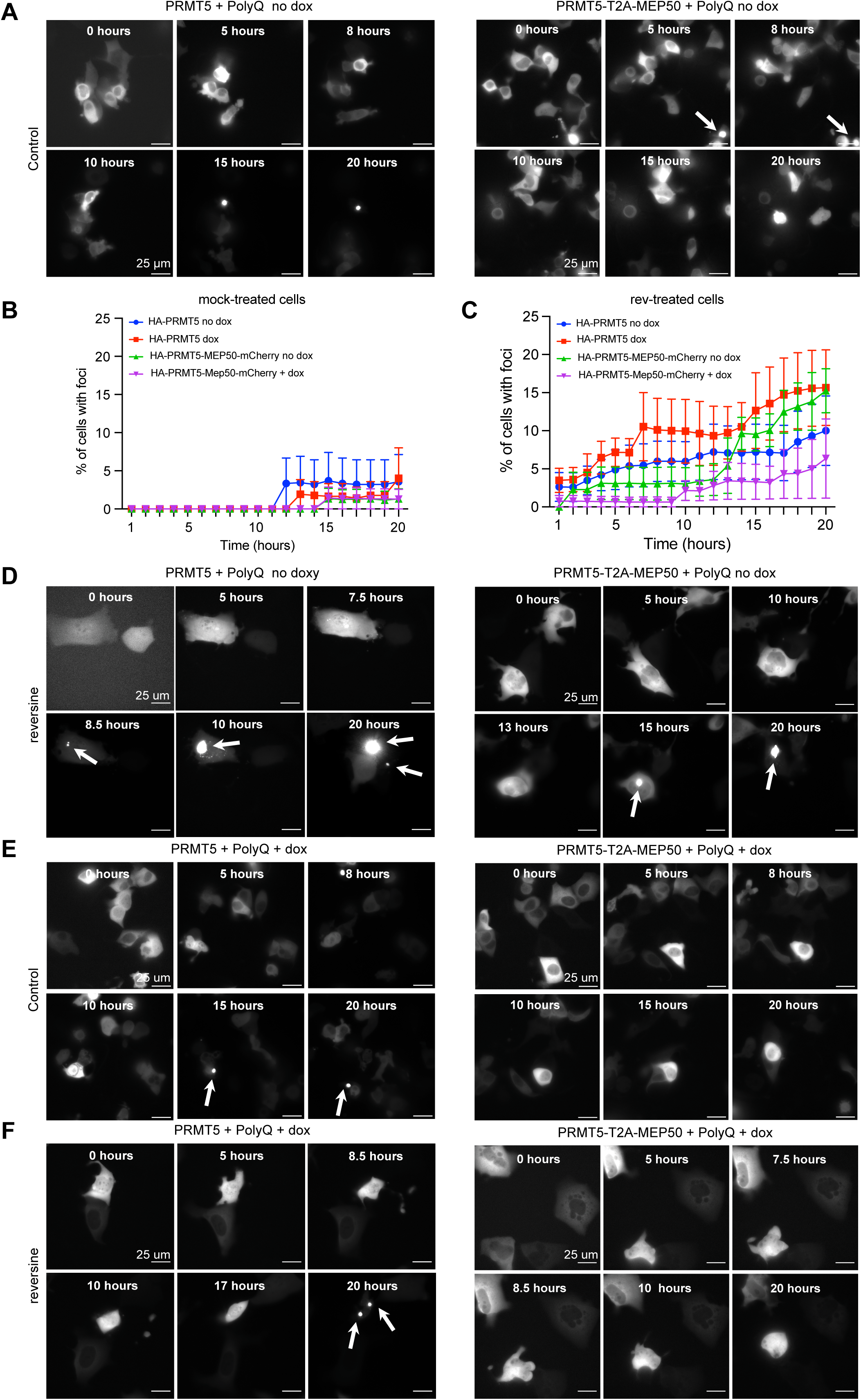
**PRMT5:MEP50 complex reduces HTT^EX^**^1^**^-Q^**^71^ **aggregation. (A)** Representative stills of live-cell imaging experiments using MCF7 HA-PRMT5 (left panels) or HA-PRMT5-T2A-MEP50-mCherry cells (right panels) transfected with HTT^EX1-Q71^-GFP, which were treated with carrier control. **(B)** Quantification of HTT^EX1-Q71^ aggregates in live-cell imaging of MCF7 HA-PRMT5 cells transfected with HTT^EX1-Q71^-GFP, treated with carrier control (blue dot), 500 nM rev treated (red square), 1000 ng/ml dox (green arrow), or 1000ng/dox plus 500nM rev (purple arrow). **(C)** Quantification of HTT^EX1-Q71^ aggregates in live-cell imaging of MCF7 HA-PRMT5-T2A-MEP50-mCherry cells transfected with HTT^EX1-Q71^-GFP and treated with carrier control (blue dot), 500 nM rev treated (red square), 1000 ng/ml dox (green arrow), and 1000ng/dox plus 500nM rev (purple arrow). Dox was added for 72h and rev for 48h prior to imaging. Data for carrier control treatments represent 2 technical replicates per condition; dox and rev treated represent 3 replicates per condition. **(D)** Representative stills of live-cell imaging experiments using MCF7 HA-PRMT5 (left panels) or HA-PRMT5-T2A-MEP50-mCherry cells (right panels) transfected with HTT^EX1-Q71^-GFP, which were rev-only treated. **(E)** Representative stills of live-cell imaging experiments using MCF7 HA-PRMT5 (left panels) or HA-PRMT5-T2A-MEP50-mCherry cells (right panels) transfected with HTT^EX1-Q71^-GFP, which were dox-treated. **(F)** Representative stills of live-cell imaging experiments using MCF7 HA-PRMT5 (left panels) or HA-PRMT5-T2A-MEP50-mCherry cells (right panels) transfected with HTT^EX1-Q71^-GFP, which were dox and rev-treated.

### Stoichiometric induction of PRMT5:MEP50 protects aneuploid cells against proteotoxic stress

We further tested our hypothesis that stoichiometrically induced PRMT5:MEP50 expression mitigates the detrimental consequences of proteotoxic stress. For this purpose, we determined proliferation of MCF7 cells with increased PRMT5 versus PRMT5:MEP50 expression under proteotoxic stress induced through the proteasome inhibitor MG132^53,54^. To quantify proliferation, cells were transduced with an H2B-mCherry construct and imaged in an IncuCyte time-lapse imager. MG132 modestly reduced cell growth rates, indicating that proteotoxic stress affects proliferation in our model (**Sup.** Fig 2A). PRMT5:MEP50 mitigated the detrimental effect of MG132 on proliferation (blue line, **Fig. 5A, B, Supp.** Fig. 2B-C), whereas PRMT5 alone did not rescue impaired proliferation in MG132 treated cells (red line, ratio dox^+^/dox^-^=∼1, **Fig. 5A**, **Supp.** Fig. 2B-C). We conclude that PRMT5:MEP50 co-expression yields a growth advantage to MCF7 breast cancer cells under proteotoxic stress. Our findings translate to other compounds and cancer cell lines, as coordinated increased expression of PRMT5 and MEP50 increases resistance to the proteasome inhibitor Oprozomib in a large panel of ∼1,600 cancer cell lines in the DepMap repository (**Fig. 5B**). In summary, stoichiometric increase in PRMT5:MEP50 expression protects cells against proteotoxic stress, whereas increased expression of PRMT5 alone does not.

**Figure 5.**
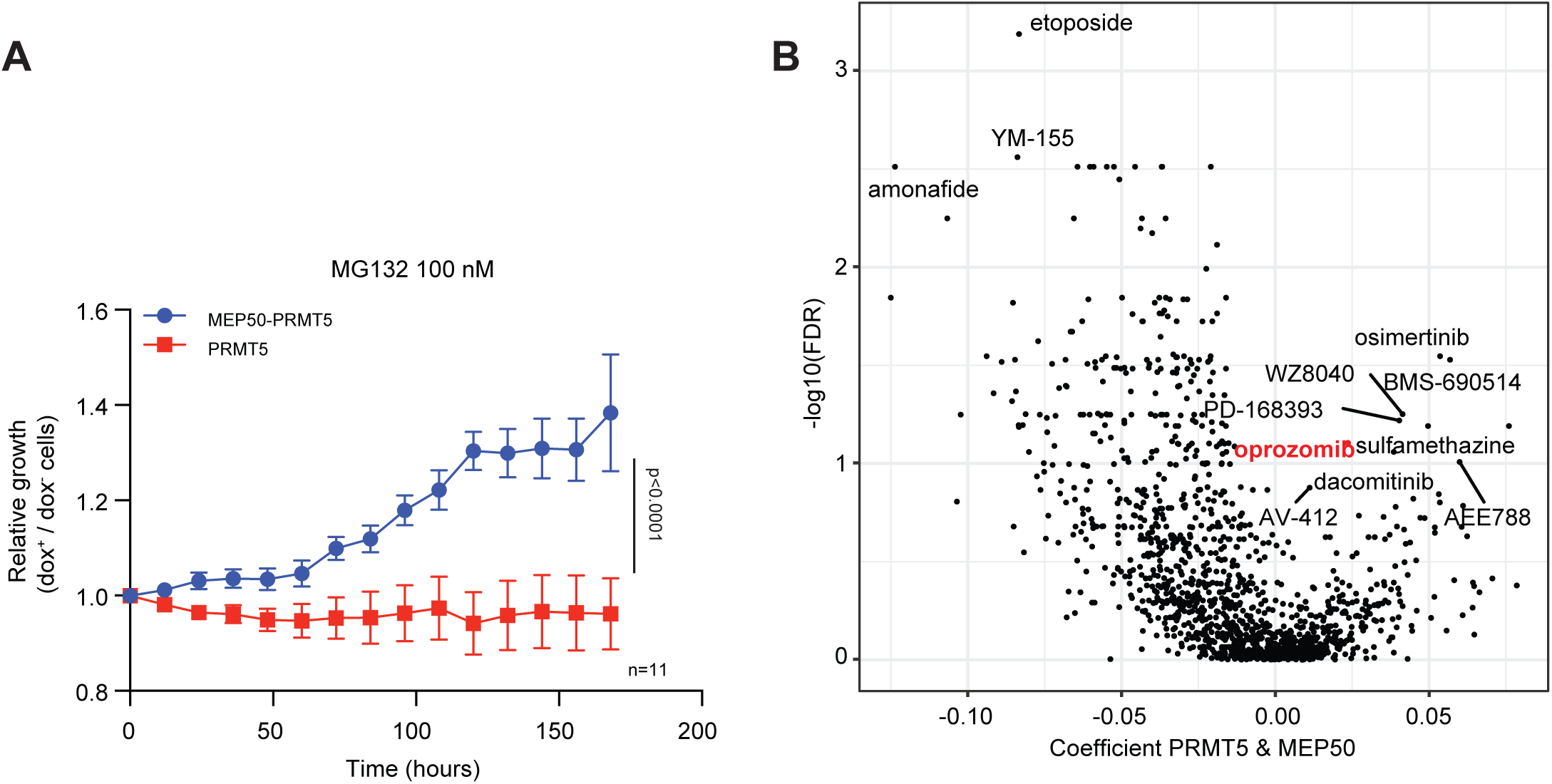
PRMT5:MEP50 expression reduces sensitivity to the proteasome inhibitor MG132. **(A)** Ratios of carrier-treated cell line normalized growth curves for dox-treated over non-dox treated HA-PRMT5 or HA-PRMT5-T2A-MEP50-mCherry MCF7 cells treated with 100 nM MG132, as determined in an IncuCyte time-lapse imager. Individual growth curves are shown in Supplemental Figure 2 A, B, C. Data are represented as mean ± SEM, p-values were calculated from an unpaired two-tailed t-test. **(B)** Volcano plot showing drug sensitivities of cancer cell lines deposited at the CCLE with high expression of PRMT5, MEP50 or PRMT5 and MEP50.

### Stoichiometric PRMT5:MEP50 co-expression and activity promote resistance to CIN

PRMT5 and MEP50 display increased expression in mouse models for CIN-driven lymphoma^22,23,55^. We therefore determined whether stoichiometrically increased PRMT5:MEP50 expression promotes resistance to CIN. We induced CIN in MCF7 cells with induced expression of PRMT5:MEP50 or PRMT5 alone using several concentrations of the MPS1 inhibitors rev or cpd-5. Cells with coordinated PRMT5:MEP50 expression were less growth-impaired by CIN than cells that only expressed PRMT5, with this effect being most evident at 250 nM rev (**Fig. 6A**, **Supp.** Fig. 2D-F) and 50 nM cpd-5 (**Fig. 6B, Supp.** Fig. 2G-I). Hence, coordinated increase in PRMT5:MEP50 expression reverses some of the toxic effects of CIN on proliferation.

**Figure 6.**
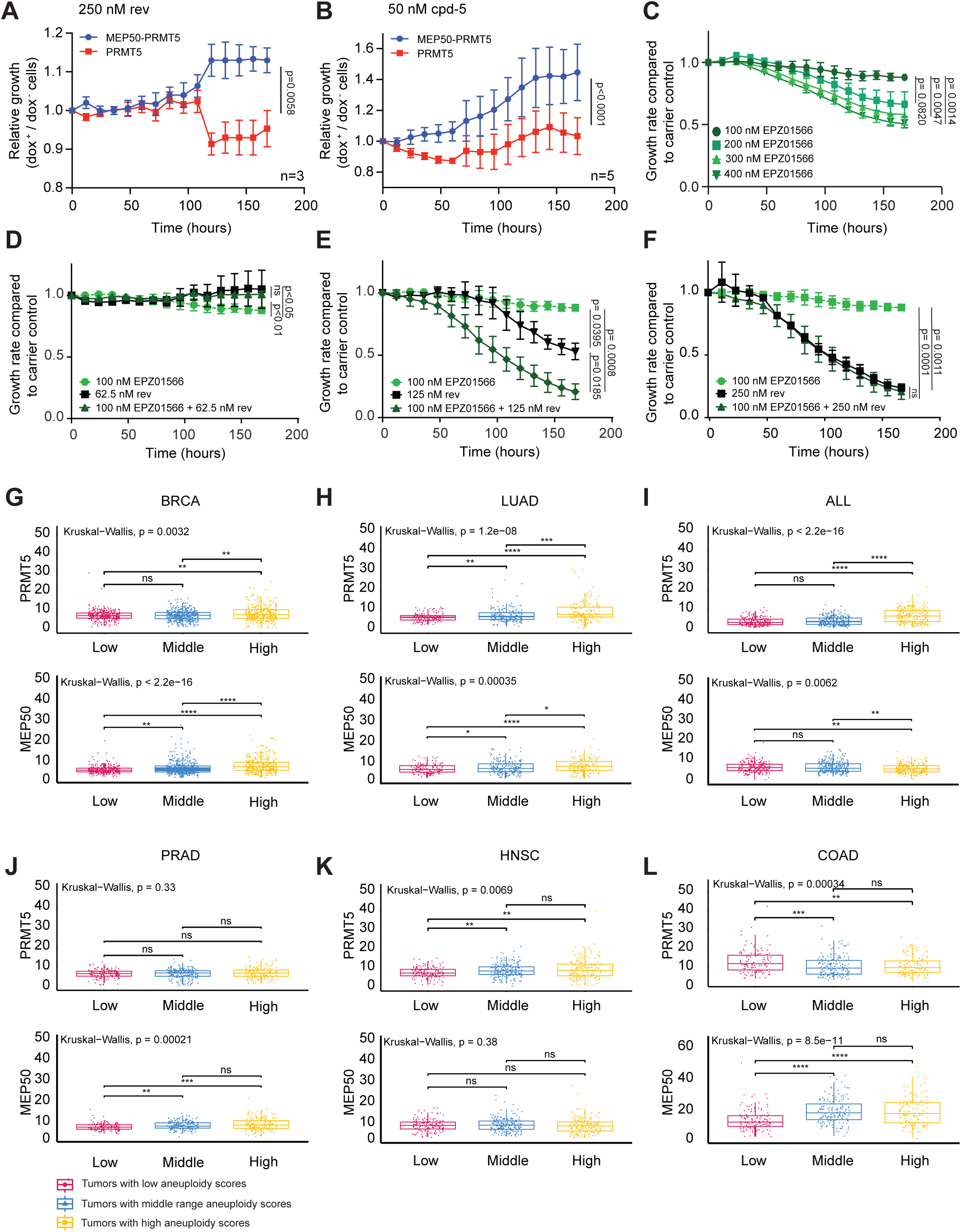
Inhibition of PRMT5:MEP50 sensitizes cells to drug-induced CIN. (A, **B)** Ratios of carrier-treated cell line normalized growth curves for dox-treated over non-dox treated HA-PRMT5 or HA-PRMT5-T2A-MEP50-mCherry MCF7 cells treated with MPS1 inhibitors, **(A)** 250 nM rev or **(B)** 50 nM cpd-5. Growth was measured in an IncuCyte time-lapse imager. Individual growth curves are shown in Supplemental Figure 2 **(D, E, F** for rev, **G, H, I** for cpd-5). Data are represented as mean ± SEM. **(C)** Ratios of carrier-treated cell line normalized growth curves of MCF7 cells treated with 100, 200, 300 and 400 nM of EPZ015666 over carrier control-treated cells. (**D, E, F**) Ratios of carrier-treated cell line normalized growth curves of MCF7 cells treated with 100 nM of EPZ015666, rev and in combination **(D)** 62.5 nM rev, **(E)** 125 nM rev or **(F)** 250 nM rev, as determined in an IncuCyte time-lapse imager. Data are represented as mean ± SEM, p-values were calculated from an unpaired two-tailed t-test. * p ≤ 0.05, ** p ≤ 0.01, ns: non-significant. **(G-L)** PRMT5 and MEP50 expression levels in TCGA-included breast cancer **(G)**, lung adenocarcinoma **(H)**, acute lymphoblastic lymphoma **(I)**, prostate adenoma **(J)**, head and neck squamous carcinoma **(K)**, and colon adenocarcinoma **(L)**, stratified for a low (red), middle range (blue) or high (yellow) aneuploidy score.

To test if PRMT5:MEP50 promotes CIN resistance via its methyltransferase activity, we exposed wild type MCF7 cells to the substrate-competitive inhibitor EPZ015666, which selectively blocks PRMT5:MEP50 activity while leaving the activity of other PRMTs intact^56,57^. Proliferation was assessed as a proxy for EPZ015666 toxicity for concentrations up to 400 nM. 100 nM EPZ015666 did not compromise proliferation and was hence considered non-toxic (**Fig. 6C**). To test if inhibition of PRMT5:MEP50 sensitizes cells to CIN, we combined a range of rev concentrations with the non-toxic dose of 100 nM EPZ015666 (**Fig. 6D-F**). 250 nM rev significantly impaired proliferation at baseline, whereas 125 nM rev was less toxic. Combining 100 nM of EPZ015666 reduced proliferation of 125 nM rev treated cells but not of cells treated with the carrier control. We conclude that inhibition of PRMT5:MEP50 activity sensitizes cells to rev, corroborating our observation that PRMT5:MEP50 protects cells against drug-induced CIN, while at the same time revealing a potential targetable vulnerability of cells with (induced) CIN.

To physiologically validate our findings, we analyzed the correlation between expression levels of PRMT5:MEP50 and aneuploidy. Both PRMT5 and MEP50 expression correlated with aneuploidy in a tissue dependent manner, with a positive correlation between an increase in expression of both PRMT5 and MEP50 and aneuploidy in the TCGA-BRCA (breast cancer), TCGA-LUAD (lung adenocarcinoma), and MP2PRT-ALL (acute lymphoblastic lymphoma) cohorts (**Fig. 6G-I**). Other cohorts such as TGCA-PRAD (prostate adenoma), TCGA-HNSC (head and neck squamous carcinoma) and TGCA-COAD (colon carcinoma) showed low or even negative correlation between PRMT5:MEP50 expression and aneuploidy (**Fig. 6J-L**). Tissue specific effects of PRMT5 in cancer have been noted earlier and are assigned, for instance, to tissue-related nuclear versus cytoplasmic distribution of PRMT5, with cancer-protective or oncogenic effects, respectively^58^.

In summary, we show that cancer cells that exhibit CIN stoichiometrically express PRMT5 and its obligatory counterpart MEP50, which is required to prevent cytoplasmic PRMT5 foci formation. Coordinated upregulation of the PRMT5:MEP50 complex reduces CIN-induced protein aggregation and sensitivity to proteotoxic stress and hence promotes resistance to CIN-inducing drugs. Conversely, chemical inhibition of PRMT5:MEP50 reduces tolerance of cancer cells to drug-induced CIN. This could be exploited as a targetable vulnerability of cells experiencing CIN-induced proteotoxic stress in cancers including breast and lung cancer and T-ALL.

## Discussion

Chromosomal instability is an enabling hallmark of cancer cells^59^ driving their evolution by promoting copy number changes that help cells adopt malignant fate. However, CIN also leads to proteotoxic, metabolic and genotoxic stress, ultimately triggering an inflammatory response^60–63^. This suggests that CIN^HIGH^ cancer cells must acquire mechanisms to cope with the detrimental consequences of CIN. Indeed, cancer cells develop compensation mechanisms to tolerate survival stresses, including ploidy-tolerating mutations^64^, protein dosage compensation^65^, whole genome doubling^66^, increased aggregation^50^, upregulated immune evasion pathways and genes^67^. In addition, aneuploidy in cancer has been linked to a worsened prognosis, aggressive disease, treatment resistance, and metastasis in patients^16,68,69^.

We previously identified *prmt5* to be amplified in a model for CIN-induced murine T-ALL^22,23^. To test if the resulting increased expression of Prmt5 helps cancer cells cope with CIN, we engineered cell models in which PRMT5 expression could be induced to the levels observed in cancer, in CIN or non-CIN backgrounds. We find that increased expression of PRMT5 leads to the formation of foci that are not SGs or PBs but appear to be protein aggregates. In line with this, and with other studies showing that CIN promotes protein aggregation^70^, induction of CIN further promotes PRMT5 foci formation. Similarly, and in agreement with previous observations^71^, CIN promotes aggregation of a PolyQ reporter protein. Comparing PRMT5 in foci to PRMT5 outside of foci revealed that free PRMT5 is bound to MEP50, a required interactor for PRMT5 substrate recognition. When we induce expression of both PRMT5 and MEP50 concomitantly, we find that accumulation of PolyQ aggregates induced by CIN is reduced. Our findings therefore suggest that increased PRMT5:MEP50 as observed in cancer contributes to aneuploidy tolerance by reducing aggregation-induced proteotoxic stress. How does PRMT5:MEP50 modulate protein aggregation? One possible mechanism is through Hsp90 methylation by PRMT5:MEP50, which has previously been reported to activate Hsp90^72^. Hsp90 is a molecular chaperone that assists in the folding and stabilization of a wide range of client proteins, including those prone to misfolding and aggregation. Its activation is predicted to reduce proteotoxic stress by enhancing the cell’s ability to manage misfolded proteins. In cancer cells, Hsp90 is often upregulated, supporting the stability of oncogenic proteins and preventing their aggregation under stress conditions^73^. PRMT5 itself has been identified as a client of Hsp90, and its stability appears to depend on Hsp90’s chaperone function^72^. Therefore, PRMT5:MEP50 may influence protein aggregation not only through direct methylation of Hsp90 but also by participating in a feedback loop that stabilizes its own activity via Hsp90. This interplay highlights a potential therapeutic vulnerability in cancers characterized by high proteotoxic stress and PRMT5-driven epigenetic dysregulation^74^.

Inhibition of PRMT5:MEP50 acted synergistically with drug-induced CIN, which might indicate a therapeutic vulnerability in cancer cells with an increased level of CIN. In line with these findings, we find that human cancer cell lines that overexpress PRMT5:MEP50 show decreased sensitivity to proteasome inhibitors, which also provoke proteotoxic stress. This suggests that high PRMT5:MEP50 levels can potentially be used as a contra-indication for proteasome inhibitor treatment, a commonly used anticancer therapy ^75^. Conversely our work also suggests that inhibition of PRMT5:MEP50 would exacerbate CIN-induced aggregation, which provides a possible intervention strategy to target aneuploid cancers. However, the pro- or antitumorigenic effect of PRMT5 depends on its localization^58^ and accordingly we find that not all cancer types show a correlation between PRTM5:MEP50 expression and aneuploidy. Further work is urgently needed to validate PRMT5 and MEP50 expression as a targetable vulnerability in aneuploid lung and breast cancer, and T-ALL, and to identify markers predictive of PRMT5’s pro- and antitumorigenic outcomes.

## Acknowledgements

We are grateful to the members of the Foijer lab for fruitful discussions. We thank Hjalmar Permentier and Marcel de Vries from the RUG/UMCG proteomics facility for assistance with mass spectrometry experiments. MFSPR is funded by a HORIZON EUROPE Marie Sklodowska-Curie postdoctoral fellowship (101068734). Gaviria Agudelo was funded by K.T acknowledges support from a Rosalind-Franklin-Fellowship of the University of Groningen, the MESI-STRAT project (Grant Agreement No 754688) which has received funding from the European Union’s Horizon 2020 research and innovation programme, and the European Partnership for the Assessment of Risks from Chemicals PARC (Grant Agreement No. 101057014) and European Research Council (ERC AdG BEYOND STRESS, Grant Agreement No. 101054429) which have received funding from the European Union’s Horizon Europe research and innovation programme. This work was supported by a Dutch Cancer Society Grant (2015-RUG-7833) to Foijer and a COLCIENCIAS fellowship (project code 727-2015) to.

## Author contributions

MFSPR, LJJ, CGA, JES, CH, AMH, P.L.B, performed experiments; MR and AvK performed bioinformatic analyses; MATMVV. contributed resources, GL and PPR conducted EPZ015666 synthesis. MSFPR, LJJ, CGA, JES, AO, KT., and FF wrote the paper; KT and FF conceptualized the study, KT and FF provided funding.

## Material and Methods

### Resource availability

#### Lead Contact

Further information and requests for resources and reagents should be directed to and will be fulfilled by the lead contact, Floris Foijer.

#### Materials Availability

Plasmids generated in this study are available from the lead contact with a completed materials transfer agreement.

#### Data and Code Availability

- All data reported in this paper are available from the lead contact upon request.
- This paper does not report original code.
- Any additional information required to reanalyze the data reported in this paper is available from the lead contact upon request.

### Experimental model details Cell lines

RPE-1 (human retinal pigmented epithelial), MCF7 (breast cancer), and HEK293T (human embryonic kidney) cells were grown in DMEM (Gibco) supplemented with 10% FBS (Sigma) and 5% Pen/Strep (Invitrogen). MCF10A (human mammary epithelial) cells were maintained in DMEM/F12 (Gibco) supplemented with 5% horse serum (Invitrogen), 20 ng/mL EGF (Peprotech), 500 ng/mL hydrocortisone (Sigma), 5% Pen/Strep (Invitrogen), 100 ng/mL cholera toxin (Sigma) and 10 µg/mL insulin (Sigma). BT549 (breast cancer) and 4T1 (breast cancer) were maintained in RPMI 1640 (Gibco) supplemented with 10% FBS (Sigma) and 5% Pen/Strep (Invitrogen). All cell lines were cultured in 5% CO_2_ in an incubator at 37°C.

### Method details

#### Plasmids

For pRetrox GFP-PRMT5, we first PCR amplified (Phusion polymerase, New England Biolabs (NEB)) GFP or mCherry with BglII and BamHI+NotI sites 5’and 3’ respectively. This PCR fragment was cloned into BamHI, and Not1 (NEB) digested pRetrox backbone (Clontech). We then PCR-amplified PRMT5 from a human PRMT5 cDNA clone (Thermo Scientific) flanked by BamHI and NotI 5’ and 3’ respectively, which was cloned into pRetrox-GFP digested with BamHI and NotI.

For pRetrox PRMT5-HA we PCR-amplified PRMT5 from pRetrox-GFP PRMT5 flanked by BglII-site, HA-tag and MluI-site. This PCR fragment was cloned in pRetrox-Tight-Pur cut with BamHI and MluI.

For pRetrox PRMT5-GFP-HA we PCR amplified EGFP from pRetrox GFP-PRMT5 flanked on both sites by XbaI sites. This PCR fragment was cloned in pRetrox PRMT5-HA cut with XbaI.

For pRetrox GFP/mCherry-Dcp1a, we PCR-amplified Dcp1a from a human Dcp1a cDNA clone (GE Healthcare) flanked by BamHI and NotI 5’ and 3’ respectively, which was cloned into pRetrox-GFP and pRetrox-mCherry digested with BamHI and NotI.

For pRetrox GFP-PRMT5-T2A-MEP50, we first PCR amplified (Phusion polymerase, New England Biolabs (NEB)) PRMT5 flanked by BamHI and NotI sites 5’ and 3’ respectively. This fragment was cloned into BamH1 and Not1 (NEB) digested pRetrox-GFP. We then PCR amplified MEP50 from human cDNA flanked by T2A-NotI and MluI sites 5’ and 3’, respectively, which was cloned into pRetrox-GFP-PRMT5 digested with NotI and MluI. For pRetrox GFP-PRMT5-T2A-MEP50-mCherry, we PCR amplified mCherry with MluI and EcoRI sites. This fragment was cloned into pRetrox GFP-PRMT5-T2A-MEP50 cut with MluI and EcoRI.

For the generation of pRetrox HA-PRMT5-T2A-MEP50-mCherry, we first PCR amplified pTight-HA flanked by BglII and BamHI sites, from a pre-existing pRetrox-HA vector. This fragment was cloned into pRetrox PRMT5-T2A-MEP50 digested with BglII and BamHI.

All primers and vectors used in this study are listed in Table S1.

#### Transfection and retroviral transduction

Retroviral particles (pRetrox system; Clontech) were produced in Turbofect (Thermo) transfected HEK293T cells. Cell lines were retrovirally transduced with pRetrox-rtTA virus and next with pRetrox GFP-PRMT5, PRMT5-GFP, PRMT5-HA, GFP-PRMT5-T2A-MEP50-mCherry, HA-PRMT5-T2A-MEP50-mCherry or GFP/mCherry-DCP1A virus.

A GFP-tagged exon1 fragment of the HTT gene containing 71 glutamines (HTT^EX1-Q71^-GFP) was kindly gifted by Ellen Nollen’s Lab from ERIBA, University Medical Centre Groningen. Transfection of MCF7 cells with the aggregation probe was performed with polyethylenimine (PEI) 24h prior imaging.

#### Western blotting

Cells were lysed in ELB (150 mM NaCl, 50 mM Hepes pH 7.5, 5 mM EDTA, 0.1 % NP-40) supplemented with Complete Protease Inhibitor (Roche). 20 µg of protein was mixed with 5x sample buffer (50% Glycerol, 10% SDS, 0,5M DTT, 250 mM Tris pH 6,8) and heated for 5 minutes at 98 °C. The samples were then separated on 10 % SDS-polyacrylamide gels and transferred onto polyvinylidene difluoride (PVDF) membranes (Millipore). After blocking in Intercept Blocking Buffer (IBB, LI-COR) 1:1 diluted with TBS (19 mM Tris base, NaCl 137 mM, KCl 2.7 mM), membranes were incubated with the primary antibodies diluted in IBB overnight at 4 °C. After 3 washes with TBS-T (19 mM Tris base, NaCl 137 mM, KCl 2.7 mM and 0.1% Tween-20), the membranes were incubated for 1 hour at RT with the fluorophore conjugated secondary antibodies diluted in IBB at room temperature. After two washes with TBS-T and one in TBS, the fluorescent signals were quantified using the Odyssey CLx Imaging System (LI-COR). Antibodies used in this study are listed in the Key Resources Table.

#### Time-lapse imaging

For time-lapse, 20000 cells per well were seeded on imaging chambers (LabTek). 48 hours prior to imaging cells were treated with 500 nM rev (Sigma) and 24 hours prior to imaging dox (Sigma) was added to induce transgene expression (concentration dox is dependent on the performed experiment, see figure legends). Cells were imaged for up to 20 hours on a DeltaVision Elite imaging station (Applied Precision, GE Healthcare). Images were taken every 4 minutes using the 40x objective. Movies were analyzed using SoftWorx suite.

For protein aggregation curves, 4000 MCF7 cells were seeded into each well of 96-well plates. Cells were treated with 1 µg/ml dox for 72 h and 500 nM rev for 48 h, and transfected with HTT^EX1-Q71^-GFP 24 h before imaging. All measurements were performed with technical duplicates as indicated. Cells were monitored by time lapse imaging using a DeltaVision Elite imaging station (Applied Precision, GE Healthcare). Images were analyzed using FIJI software.

#### Z-stack imagse acquisition

Z-stack images to examine the co-localization between PRMT5 and DCP1a were acquired using Z-stack mode on Zeiss LSM 780 NLO microscope (PlanNeofluar 63x/1.31mm CorrDIC glycerine immersion). Cells grown on coverslips were placed in the microscope imaging chamber. A 488 nm laser was used to acquire GFP-tagged PRMT5 images and mCherry-tagged DCP1a images were obtained using a 561 nm laser. The images obtained were processed using Imaris software.

#### Immunofluorescence

RPE-1 and MCF10A cells were grown in 6-well plates, with each well containing a sterile glass cover slip (VWR international). To induce stress granules, cells were treated with 500 µM arsenite (Sigma) for 60 minutes. Cells were fixed in 4% paraformaldehyde in PBS for 5 minutes at room temperature. After 3 washing steps with PBS the cells were permeabilized with 1 mL 0.1% Triton-100 (Sigma) for 1 minute at room temperature. After two washes with PBS cells were blocked by incubating the cells in PBS with 3% FCS for 20 minutes at room temperature. This was followed by an overnight incubation with 200 µL of the primary antibodies (see table 3) diluted in blocking buffer at 4°C. After 3 washes with PBS, cells were incubated with secondary antibodies (see table 3) diluted in blocking buffer for 30 minutes in a humid dark chamber at room temperature. Finally, cells were washed for 3 times with PBS and 2 times with water. Finally, cells were mounted with 30 µL Vectashield mounting medium with DAPI (Vector laboratories). Immunofluorescence microscopy was performed using a Zeiss observer and Apoptome.2 or a DeltaVision Elite imaging station (Applied Precision, GE Healthcare). Images were analyzed using the ZEN2012 blue software or SoftWorx suite. Granule intensity was quantified using ImageJ. Antibodies used in this study are listed in Key Resources Table.

#### FRAP analysis

FRAP experiments were performed using 3D-FRAP system built on Zeiss LSM 780 NLO microscope (PlanNeofluar 63x/1.3Imm CorrDIC glycerine immersion). Similar to the Z-stack experiments, cells grown on coverslips were placed in the microsope imaging chamber. Experiments were performed at 37 °C with 5% CO2 using a live cell chamber system (Incubator XL S1 DARK). For each acquisition, PRMT5-GFP or GFP-PRMT5 granules were bleached using a 488 nm laser, while mCherry-DCP1a granules were bleached with a 561 nm laser. Ten pre-bleach images were acquired. Post-bleach images were acquired every 2 seconds. The experiments were analyzed using ZEN acquisition software.

#### Isolation of proteins in foci

RPE-1 GFP-PRMT5/PRMT5-HA-GFP cells were treated with dox (Sigma) for 3 days. The cells were washed with cold PBS 3 times and lysed with elution buffer (150 mM NaCl, 50 mM HEPES pH 7.5, 5mM EDTA, 0.1% NP-40) supplemented with Complete Protease Inhibitor (Roche) and 1 mM DTT. Cells were collected and fresh elution buffer was added after centrifugation (4°C, 17000 x g, 10 minutes). The supernatant is depicted as ‘non-granule’ fraction in the figures. The pellet was resuspended in elution buffer and passed through a 20G needle and mixed 3 times with a homogenizer (Polytron ®) (15000 rpm, 1 minute) with cooling on ice in between. After centrifugation (4 °C, 600 x g, 10 minutes) the supernatant was taken for another centrifugation step (4 °C, 17.000 x g, 10 minutes). The pellet containing the PRMT5 granules was resuspended in elution buffer. The PRMT5 granules were isolated from the samples by sorting for GFP using fluorescence activated cell sorting or by immunoprecipitation (described in the next section). These samples are described in the figures as “PRMT5 foci”.

#### Immunoprecipitation

For immunoprecipitation either 5 µL anti-HA (Pierce) or anti-GFP (Chromotek) beads were pre-washed with 175 µL 0.05% TBS-T and gently vortexed. The beads were collected by placing the Eppendorf tube with the sample into a magnetic stand. The supernatant was discarded, and 1 mL TBS-T was added. After gently vortexing the tube was placed back into a magnetic stand and again supernatant was discarded. The pre-washed magnetic beads were rotated while incubating for 1 hour at 4 °C with 200 µL sample containing HA or GFP-tagged PRMT5. The beads were collected with a magnetic stand and supernatant was saved (unbound sample). Beads were mixed and washed twice with 300 µL TBS-T. To elute the protein from the beads 50 µL NaOH (50mM) was added to the sample. After gently vortexing, the samples were incubated at room temperature for 10 minutes. Then the beads were magnetically separated and the supernatant containing the target antigen was saved. The samples were neutralized by adding 2.5 µL 1M HCl and sent to the mass spec facility for further analysis.

#### Mass spectrometry

Proteins were fractionated by SDS-PAGE at 60V for 6 minutes for a total migration length of ∼0.5 cm. Gels were rinsed in deionized water and stained overnight with Coomassie G250 (BioRad). Proteins in the entire 0.5 cm gel section underwent standard in-gel tryptic digestion including reduction and alkylation. Peptides were extracted from each gel section and fractionated by a nanoflow reversed-phase ultra-high pressure liquid chromatography system (nanoLC, Dionex) in-line with a Q-Exactive plus mass spectrometer (Thermo Scientific). Samples were analyzed with a 1.5 h linear gradient (3– 50% acetonitrile with 0.1% formic acid), and data were acquired in a data-dependent manner using a top 12 method with a dynamic exclusion of 20 seconds.

#### Mass spectrometry data processing

The software PEAKS 8.5 (Bioinformatics Solutions Inc., Waterloo, Ontario, Canada) was applied to the spectra generated by a Q-Exactive plus mass spectrometer to search against a Human Protein database (trEmble/SwissProt entries) (Uniprot). The identified peptides were converted into gene id’s, which were used for gene ontology analysis using Enrichr^70^.

#### IncuCyte growth curves

4000 MCF7 cells were seeded into each well of 96-well plates. Cells were treated for 72h with 1 μg/ml dox and drugs as indicated immediately before imaging. Cell growth was monitored every 12 h using an IncuCyte Zoom live-cell analysis system (Essen BioScience Ltd.). Drug-containing media were refreshed at day 4. Cell confluency was normalized to seeding density and control-treated cells as indicated in the text.

#### Quantification and statistical analyses

Protein bands were quantified using the Image Studio Lite Software (LI-COR). Image J was used to quantify the fluorescence intensity of the images. Cell density was quantified using IncuCyte ZOOM 2019B software Gene ontology analysis was performed with Enrichr. Graph plots and statistical analysis were performed in Graphpad Prism 9 software according to the tests described in the figure legends and are shown as mean ± S.E.M. The number of experimental replicates and statistical significance (p value) are indicated in the figure legends. *, p ≤ 0.05; **, p ≤ 0.01, *** p ≤ 0.001, and n.s. represents “not significant”.

#### TCGA data analysis

Data was accessed from the GDC between the 18th and 26th of August, 2025. Samples were filtered to ensure matched measurements of bulk RNA and DNA were present for the selected patient. Only primary samples, not metastatic samples were considered for analysis. PRMT5 and MEP50 expression was defined as the Fragments Per Kilobase of transcript per Million mapped fragments (FPKM). Aneuploidy was calculated according to the following formula:

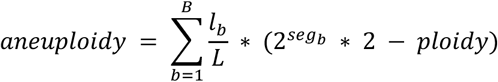

Where lb is the length of a bin, L the total length of all bins, segb the segment mean of bin b and ploidy is the base ploidy of a normal cell – in this case 2. This formula effectively weighs the contribution of a bin to the total aneuploidy score by its size.

---

**Supplemental Figure 1.**
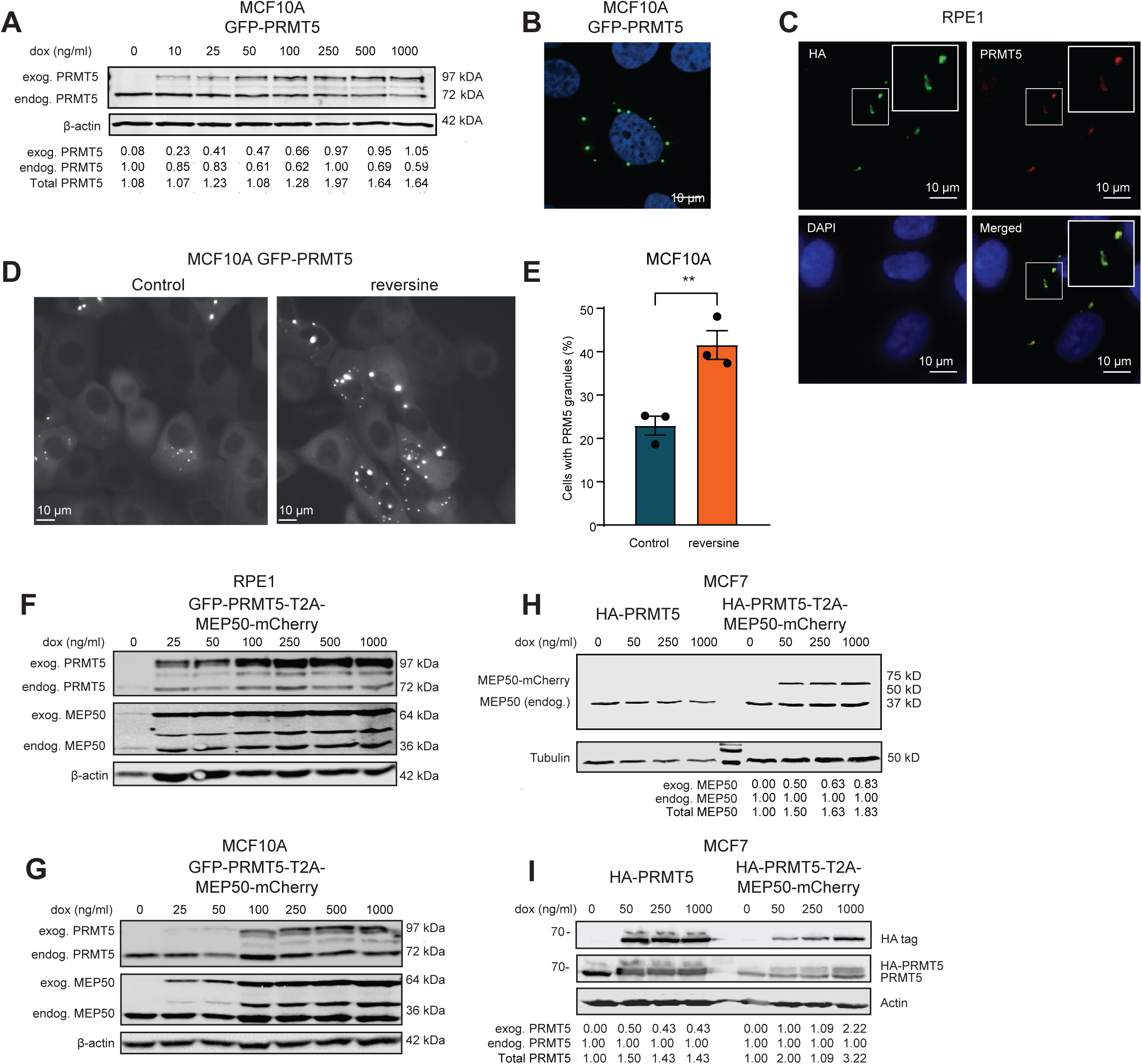
Overexpressed PRMT5 accumulates in cytoplasmic foci and stoichiometric imbalances of the PRMT5-MEP50 complex lead to PRMT5 aggregation. **(A)** Immunoblots showing PRMT5 and β-actin expression in MCF10A rtTA GFP-PRMT5 cells treated for 48 h with 0-1000 ng/ml dox. Quantification of total PRMT5 expression normalized to endogenous PRMT5 without dox is shown at the bottom of the blot. (**B**) Detection of GFP-tagged PRMT5 and nuclei/DAPI by immunofluorescence microscopy in MCF10A rtTA GFP-PRMT5 cells treated for 24 h with 500 ng/ml dox. (**C**) Detection of HA-tagged PRMT5 by immunofluorescence microscopy in RPE1 rtTA PRMT5-HA cells treated for 24h with 100 ng/ml dox using anti-HA and ati-PRMT5 antibodies. Scale bar = 10 μm. (D) Representative stills of live-cell imaging of MCF10A rtTA GFP-PRMT5 treated for 48h with 500 nM rev and 24h with 500 ng/ml dox prior to imaging. Scale bar is 10 μm. **(E)** Quantification of live cell imaging shown in **(D)** After 17 hours, at least 60 cells were counted. For statistical analysis, a two-tailed student’s t-test was applied. Data represent 3 biological replicates. **(F-I)** Immunoblot showing PRMT5, MEP50 and β-actin/Tubulin expression in **(F)** RPE1 rtTA GFP-PRMT5-T2A-MEP50-mCherry, **(G)** MCF10A rtTA GFP-PRMT5-T2A-MEP50, **(H)** MCF7 rtTA HA-PRMT5, **(I)** MCF7 rtTA HA-PRMT5-T2A-MEP50-mCherry cells. Cells were treated for 48 h with 0-1000 ng/ml dox. **(H, I)** Quantification of total MEP50 expression normalized to endogenous MEP50 without dox is shown underneath the blot panels.

**Supplemental Figure 2.**
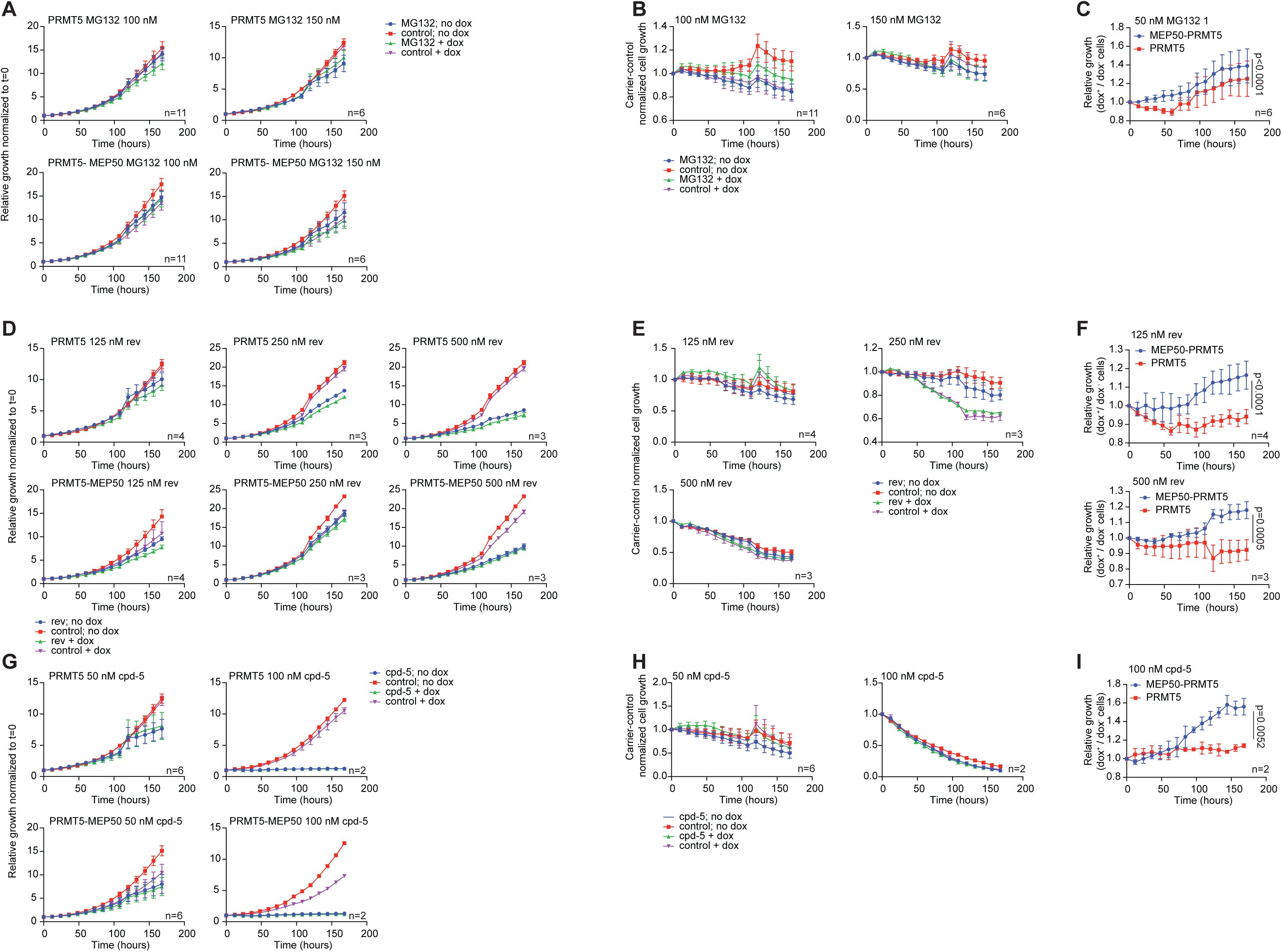
PRMT5:MEP50 reduces sensitivity to MPS1 inhibitors and the proteasome inhibitor MG132. **(A)** Growth curves normalized for seeding density of dox inducible PRMT5 or PRMT5-MEP50 overexpressing cell lines treated with various concentrations of the proteasome inhibitor MG132 in absence or presence of dox. **(B)** Carrier-control cell line normalized for dox inducible PRMT5 or PRMT5-MEP50 overexpressing cell lines induced or not with dox and treated with various concentrations of the proteasome inhibitor MG132. **(C)** Ratios between curves shown in (B) to compare effects observed between cells expressing PRMT5:MEP50 to cells expressing PRMT5 alone. Growth curves were determined with an IncuCyte time lapse imager. Error bars represent the SEM of indicated number (n) of biological replicates. p-values were calculated from an unpaired two-tailed t-test. **(D)** Seeding density-normalized growth curves for dox inducible PRMT5 or PRMT5-MEP50 overexpressing cell lines induced or not with dox and treated with various concentrations of rev. **(E)** DMSO-control cell line normalized for dox inducible PRMT5 or PRMT5-MEP50 overexpressing cell lines induced or not with dox and treated with various concentrations of the MPS1 inhibitors rev. **(F)** Ratios of DMSO-treated cell-line normalized growth curves for dox-treated over non-dox treated HA-PRMT5 or HA-PRMT5-T2A-MEP50-mCherry MCF7 cell treated with various concentrations of the MPS1 inhibitors rev. Growth curves were determined with an IncuCyte time lapse imager. Error bars represent the SEM of indicated number (n) of biological replicates. p-values were calculated from an unpaired two-tailed t-test. **(G)** Seeding density-normalized growth curves for dox inducible PRMT5 or PRMT5-MEP50 overexpressing cell lines induced or not with dox and treated with various concentrations of MPS1 inhibitor cpd-5. **(H)** DMSO-control cell line normalized for dox inducible PRMT5 or PRMT5-MEP50 overexpressing cell lines induced or not with dox and treated with various concentrations of the MPS1 inhibitor cpd-5. **(I)** Ratios of DMSO-treated cell-line normalized growth curves for dox-treated over non-dox treated HA-PRMT5 or HA-PRMT5-T2A-MEP50-mCherry MCF7 cell treated with 100nM s of the MPS1 inhibitor cpd-5. Growth curves were determined with an IncuCyte time lapse imager. Error bars represent the SEM of indicated number (n) of biological replicates. p-values were calculated from an unpaired two-tailed t-test.

## Supplemental tables

**Table S1.**
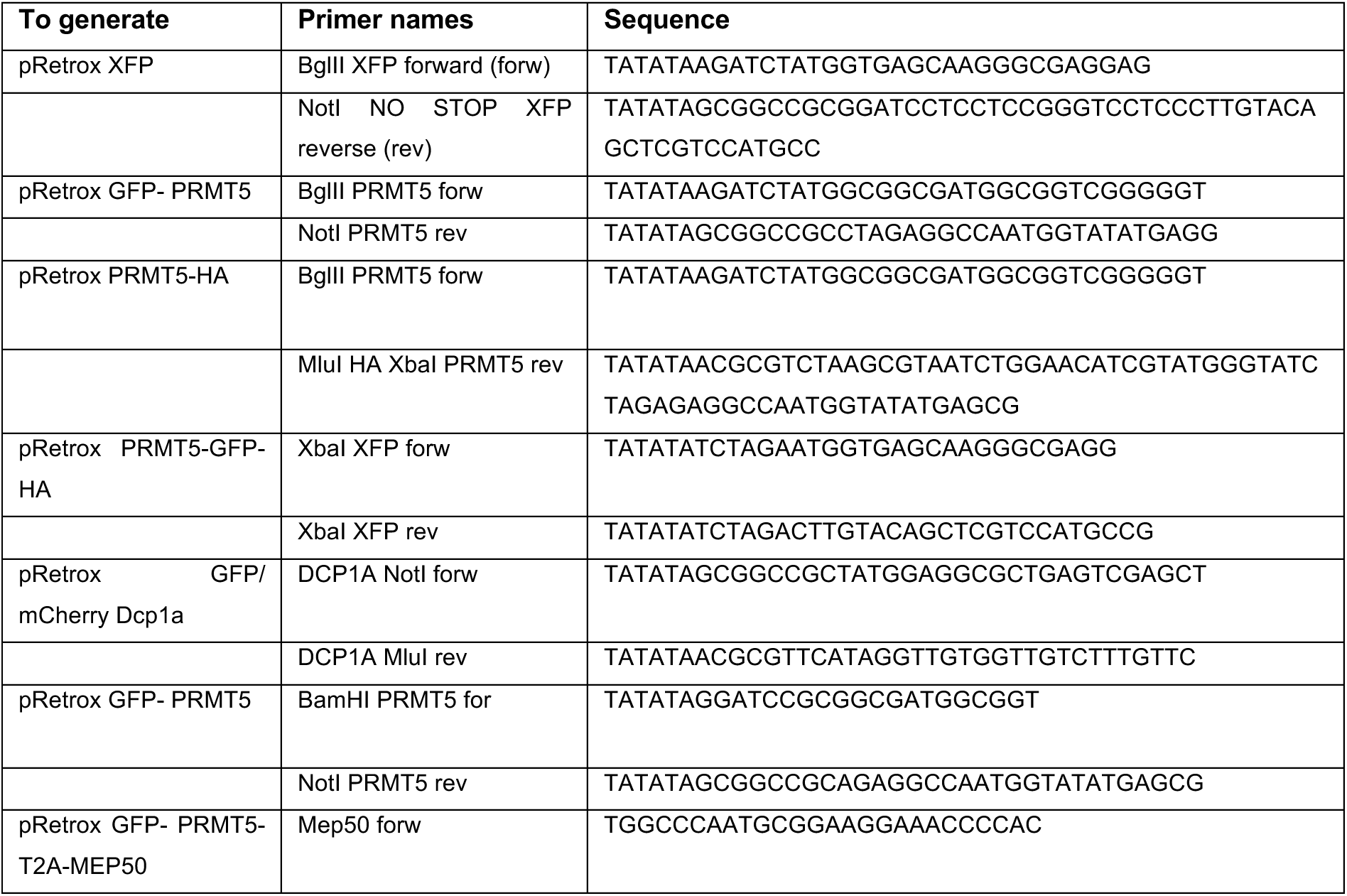
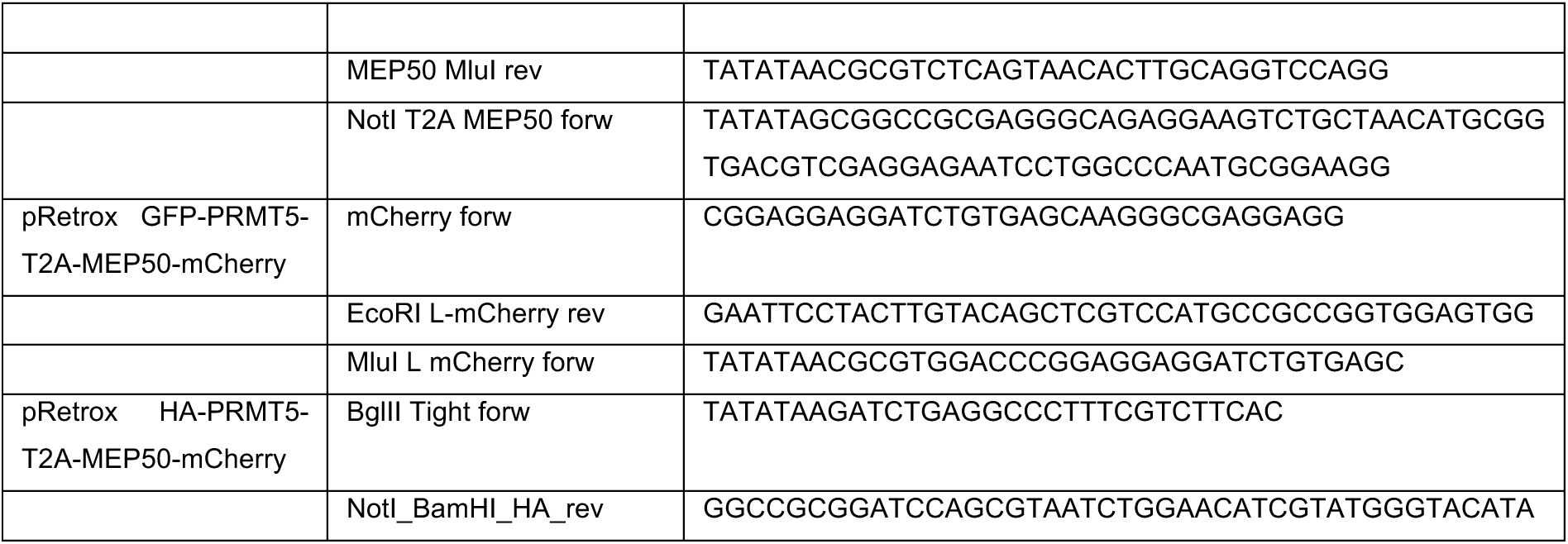
List of primers, related to STAR METHODS: Key Resources Table.

**Table S2.** Overview mass spectrometry samples, related to Figure 3.

| PRMT5 not in foci | PRMT5 foci - control | PRMT5 foci - rev |
| --- | --- | --- |
| IP_PRMT5-GFP_dox100ng/ml | IP_PRMT5-HA_dox100ng/ml | FACS_PRMT5-GFP_rev<br>treated_dox15ng/ml |
| IP_PRMT5-HA_dox15ng/ml | IP_GFP-PRMT5_dox100ng/ml |  |
| IP_PRMT5-HA_dox100ng/ml | FACS_PRMT5-GFP_dox15ng/ml |  |
| IP_GFP-PRMT5_dox100ng/ml |  |  |
| IP_PRMT5-HA_Rev<br>treated_dox15ng/ml |  |  |

